# CAR T cell engineering impacts antigen-independent activation and co-inhibition

**DOI:** 10.1101/2025.01.20.631849

**Authors:** Simon Stücheli, Christoph Schultheiß, Paul Schmidt-Barbo, Andreas Zingg, Natascha Franz, Sarah Adamo, Claudia Fischer, Heinz Läubli, Mascha Binder

## Abstract

Viral vectors have successfully modified T cells to express chimeric antigen receptors (CAR), leading to clinical approvals. However, their high cost and regulatory challenges hinder rapid and broad clinical translation. Here, we demonstrate that our lentivirally (LV) manufactured R110-CAR T cells, targeting a leukemia neoepitope, can also be engineered using non-viral Sleeping Beauty (SB) transposition with minimal-sized DNA vectors. Flow cytometry and single-cell sequencing was used to compare the two production modes using healthy donor and CLL patient-derived T cells and a CD19-CAR T cell control. SB products were shifted towards CD8^+^ subsets with expression of activation/co-inhibition markers (CD69, LAG-3, TIM-3) despite their naïve-like phenotype and lack of antigenic challenge. The CAR binding moiety modulated these patterns with R110-CAR T cells showing more aberrant phenotypes. Moreover, SB engineering resulted in inflammatory signatures along with RIG-I-like and TOLL-like nucleotide sensing potentially resulting from the transfection procedure. Patient-derived products showed significantly fewer CAR-expressing cells, reduced proliferation clusters, and lower T cell diversity, particularly with SB manufacturing, pointing at potential challenges with this method when engineering CLL T cells. Together, our data suggest that the engineering mode may substantially influence T cell properties and that these are further modulated by the CAR binding moiety and the type of T cell donor.

## Introduction

Chimeric antigen receptor (CAR) T cells are powerful tools in oncology, offering a promising approach for targeted cancer therapy (1, 2). However, beyond the availability of tumor-specific target molecules, logistical complexity and high costs of manufacturing of autologous viral products limit CAR T cell availability (3, 4). Other manufacturing procedures such as sleeping beauty (SB) transposition of CAR genes may overcome some of these – especially financial – limitations, but such products are still experimental and no commercial SB-manufactured product has been licenced so far (5, 6).

Comparative studies on CAR T cell properties according to the mode of production are scarce, especially regarding their activation and co-inhibition profile. In the context of autologous CAR T cell therapy, where a patient’s own often dysfunctional T cells are harvested, engineered, and reinfused, T cell inhibition is particularly problematic (7). When these T cells are further subjected to the stress of genetic modification and expansion, their efficacy can be further diminished. This is exemplified by CAR T cells from healthy donors being better at controling tumor growth in an in vivo model compared to CAR T cells derived from chronic lymphocytic leukemia (CLL) patients (8). Moreover, certain single-chain variable fragments (scFvs) are associated with ligand-independent tonic signalling, which has been shown to reduce the efficacy of CAR T cells, thereby limiting their therapeutic potential (9–11). This includes diminished cytokine production, reduced proliferative capacity, and impaired cytotoxic activity accompanied by the upregulation of inhibitory receptors such as PD-1, TIM-3, and LAG-3 (9, 12). Whether the CAR-inherent level of tonic signaling may be modulated by the production mode remains unclear at this point.

In the work presented here, we show that our previously published lentivirally (LV) manufactured CAR T cells that are directed against the CLL neoepitope IGLV3-21^R110^ (named anti-R110) (13), can be engineered through non-viral SB transposition of CAR genes. This production method resulted in about equal efficacy of transgene expression, but the pattern of T cell differentiation, activation and co-inhibition markers were substantially different and partially modulated by the specific CAR.

## Materials and Methods

### Cell lines and CLL patient and healthy donor blood cells

HEK293T, K562 and NALM-6 cells were purchased from the DSMZ (German Collection of Microorganisms and Cell Cultures GmbH). For ectopic expression of the IGLV3-21^R110^ light chain in NALM-6 cells (NALM-6-R110), the coding sequence was cloned into the Lentiviral Gene Ontology (LeGO) vector LeGO-iZeo2 and positively transfected cells were selected by culturing cells in the presence of 0.3 mg/mL zeocin (#ant-zn-1, InvivoGen, Toulouse, France). NALM-6 and NALM-6-R110 cells were maintained in RPMI-1640 (#61870, Thermo Fisher Scientific, Waltham, MA, USA), K562 in IMDM (#I3390, Sigma-Aldrich, St. Louis, MO, USA) and HEK293T in DMEM (#31966-021, Thermo Fisher Scientific) medium. All three media were supplemented with 10% heat-inactivated fetal bovine serum, 100 U/mL penicillin and 100 μg/mL streptomycin.

Blood samples from CLL patients were collected after informed consent as approved by the ethics committees of Halle-Wittenberg. Peripheral mononuclear cells (PBMCs) from CLL patients or healthy donors were isolated by Ficoll gradient centrifugation and primary T cells isolated using the Pan T Cell Isolation Kit (#130-096-535, Miltenyi Biotec, Bergisch Gladbach, Germany).

### Lentiviral production

Five million HEK293T cells were seeded per 10 cm dish. After 24 hours, the medium was replaced with fresh medium containing 25 μM chloroquine. The cells were then transfected with the lentiviral vector containing the CAR sequence of interest along with lentiviral packaging plasmids using the calcium phosphate transfection method (13). Five hours post-transfection, the medium was changed and the cells were incubated for three days. Subsequently, the supernatant was collected, filtered (0.2 μm), mixed with a 40% w/v PEG-8000 solution (25% PEG solution, 75% supernatant) and stored at 4°C overnight. The mixture was centrifuged (4°C, 45 minutes, 1500 rcf), the supernatant discarded, and the viral particles resuspended in activation medium (RPMI-1640, 10% heat-inactivated human serum, 100 U/mL penicillin, 100 μg/mL streptomycin, 50 μM 2-mercaptoethanol, 25 IU/mL IL-2 [#78036.1, StemCell Technologies, Vancouver, BC, Canada], 10 ng/mL IL-15 [#200-15-50UG, PeproTech, Thermo Fisher Scientific]). The viral particles were either immediately used for T cell transfection or stored at −80°C.

### Preparation of CAR T cells by lentiviral transduction or sleeping beauty transposition

For lentiviral transduction, T cells were activated using CD3/CD28 Dynabeads (#11132D, Thermo-Fisher Scientific) in a 24-well format (1 × 10L cells per well, bead-to-cell ratio 1:1) in activation medium. After 24 hours, 66% of the medium was replaced with fresh activation medium containing 0.5 mg/mL Synperonic-F108 and lentivirus encoding the respective CAR constructs. Following an additional 24 hours, 50% of the medium was replaced with fresh activation medium. After another 48 hours, the T cells were transferred to PRIME XV T-Cell CDM (91154-1L, FUJIFILM Irvine Scientific, Santa Ana, CA, USA) containing 10 ng/mL IL-15, and placed in a G-Rex-24-well format (#80192M, Wilson Wolf, St. Paul, MN, USA). On day 6 (144 hours post-T cell seeding), the CD3/CD28 beads were removed, and the medium was replaced.

For CAR T cell production using the SB transposase system, T cells were similarly activated using CD3/CD28 Dynabeads (bead-to-cell ratio 1:1) in activation medium. After 72 hours, 10 × 10L T cells were electroporated using the MaxCyte GT Electroporation System (MaxCyte, Rockville, MD, USA) following its T Cell Protocol 3. The electroporation was performed in the presence of 40 μg/mL SB transposase mRNA (custom-made by Etherna [Niel, Belgium] with CleanCap [TriLink, San Diego, CA, USA] and a plasmid-encoded polyA tail [94A]) and 150 μg/mL GenCircle CAR vector construct (GenScript, Piscataway, NJ, USA) in PRIME XV T-Cell CDM, using R-50×3 cuvettes (#ER050U3-10, MaxCyte). Electroporated T cells were immediately transferred to a G-Rex-24-well format, and after four hours, IL-15 was added to achieve a final concentration of 10 ng/mL. On day 6 (144 hours post-T cell seeding), the CD3/CD28 beads were removed.

All CAR T cells used in this study were generated from freshly isolated T cells and cultured at 37°C in a 5% CO_2_ atmosphere in PRIME XV T-Cell CDM with 10 ng/mL IL-15. The medium was refreshed once per week.

### In vitro coculture assays

Target cells seeded at 3 × 10^4^ cells/well in a 96-well plate were co-incubated with effector cells at effector-to-target (E:T) ratio 5:1 or 1:1 in standard RPMI-1640 media (10% heat-inactivated fetale bovine serum, 100 U/mL penicillin, 100 μg/mL streptomycin). Only scFv^+^ CAR T cells as determined by flow cytometry were considered effector cells. Immediately after combining target and effector cells, half of each well was subjected to flow cytometric analysis and the other half incubated for 24 hours.

Killing of target cells was assessed by gating for viable (DAPI-negative, #130-111-570, Miltenyi Biotec) CD3^-^,CD19^+^,R110^+^ target NALM-6-R110 cells and by normalizing their percentage after 24 hour incubation to 0 hour incubation time. The same normalization was applied to DAPI-negative,CD3^-^,CD19^-^,R110^-^ non-target K562 cells. Furthermore, the resulting values were used for an additional normalization step (Equation 1).

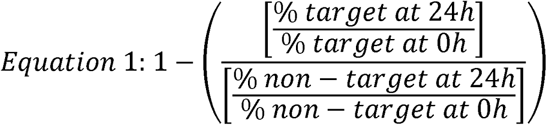

Flow cytometry was performed by washing the cells with PBS, followed by an incubation with primary unlabeled antibodies in staining buffer (1% w/v bovine serum albumin in PBS) for 30 min at 4°C in the dark. Cells were washed washed with PBS, incubated with labeled secondary antibodies and labeled primary antibodies in staining buffer for 30 min at 4°C in the dark, washed by PBS, resuspended in DAPI (1:200 dilution) and measured on a CytoFLEX S (Beckman Coulter, Brea, CA, USA). T-distributed stochastic neighbor embedding (tSNE (14)) analysis was performed by concatenating the duplicate measurements of the samples and using the embedded tSNE function of FlowJo (v10.10) with standard settings after gating for single DAPI^-^,CD3^+^ cells. Antibodies used for flow cytometry were diluted at 1:200 unless otherwise specified and listed in Supplementary Table 1.

### Single-cell RNA and VDJ sequencing

Single-cell RNA and T cell receptor (TCR) VDJ sequencing was performed using the Chromium Next GEM Single Cell 5’ Kit v2, Human TCR Amplification, and Human TCR Amplification kits (10X Genomics, Pleasanton, CA, United States) as described previously (15, 16). For library preparation, live (DAPI-negative) T cells from the SB and LV expansion cultures were FACS-sorted after staining the cells at 4°C for 30 min using a FITC-labeled anti-CD3 (clone SK7, #345763, BD BioSciences, Franklin Lakes, NJ, USA) antibody. Quality control steps were performed using an Agilent TapeStation (Agilent Technologies, Inc., Santa Clara, CA, United States). Sequencing was done on an Illumina NovaSeq 6000 system (Illumina, San Diego, CA, United States). Raw gene expression and TCR data were processed to read counts with the Cell Ranger pipeline (v7.0.1) (10X Genomics), which includes alignment to the human reference genome (GRch38), filtering and barcode counting. Processed read count matrices were analyzed and plotted using Seurat (v5.0.3) (17), scRepertoire (18) or scplotter package (https://github.com/pwwang/scplotter) in RStudio (v2023.06.1; R version 4.3.1 (2023-06- 16)). Integration of the single samples were performed using Harmony (19). To detect CAR T cell-specific sequences within the data, we created a custom reference genome based on the applied CAR T construct sequences using the cellranger mkref pipeline. Mean cluster expression per feature was calculated using Seurat’s *AggregateExpression* function. Gene set scores were calculated and plotted using the UCell package (20). Heatmaps were generated using the pheatmap package. Differentially expressed genes (DEGs) were identified using Seurat’s *FindMarkers* function (test.use = “wilcox”, min.pct = 0.1, logfc.threshold = 0.1) and visualized using the EnhancedVolcano package. Gene set enrichment analyses (GSEA) were performed using the fgsea package (version 1.26.0). Human Hallmark category gene sets (e. g. KEGG, GO) were obtained from the Molecular Signatures Database (MSigDB) using the msigdbr R package. DEGs were ranked based on their average log2 fold change and the resulting gene list served as input for the GSEA analysis.

## Results

### Engineering and expansion of CD19- and R110-CAR T cells by LV transduction versus SB transposition

We used two constructs with potential applicability in CLL: An anti-CD19 CAR sequence containing 4-1BB and CD3ζ and a CAR sequence against the CLL neoantigen IGLV3- 21^R110^ containing the same elements (Figure 1A). Both sequences were used to construct human T cells expressing the respective CAR either by LV transduction or by SB transposition (Figure 1B). We compared LV- and SB-manufactured CAR T cells derived from healthy donor blood. Despite the cell damage caused by the electroporation procedure for the SB-engineered products, T cell expansion and cell viability recovered within the first week. Figure 1C shows the gating scheme for CAR-expressing T cells using the scFv-linker as detection marker. The CAR was expressed by approximately 40% of transduced or electroporated T cells two weeks from manipulation (Figure 1D). With SB transposition, we achieved somewhat lower percentages of CAR expression when CLL-derived T cells were used, potentially due to their greater fragility (Figure 1D). scFv membrane density was estimated based on flow cytometry measured mean fluorescence intensity (MFI) of the CAR linker. After stable integration, scFv densities showed no consistent differences between SB- versus LV-manufactured CAR T cells (Figure 1E). R110-CAR expression was, however, consistently lower compared to CD19-CAR expression in both SB- and LV-manufactured products (Figure 1E).

**Figure 1.**
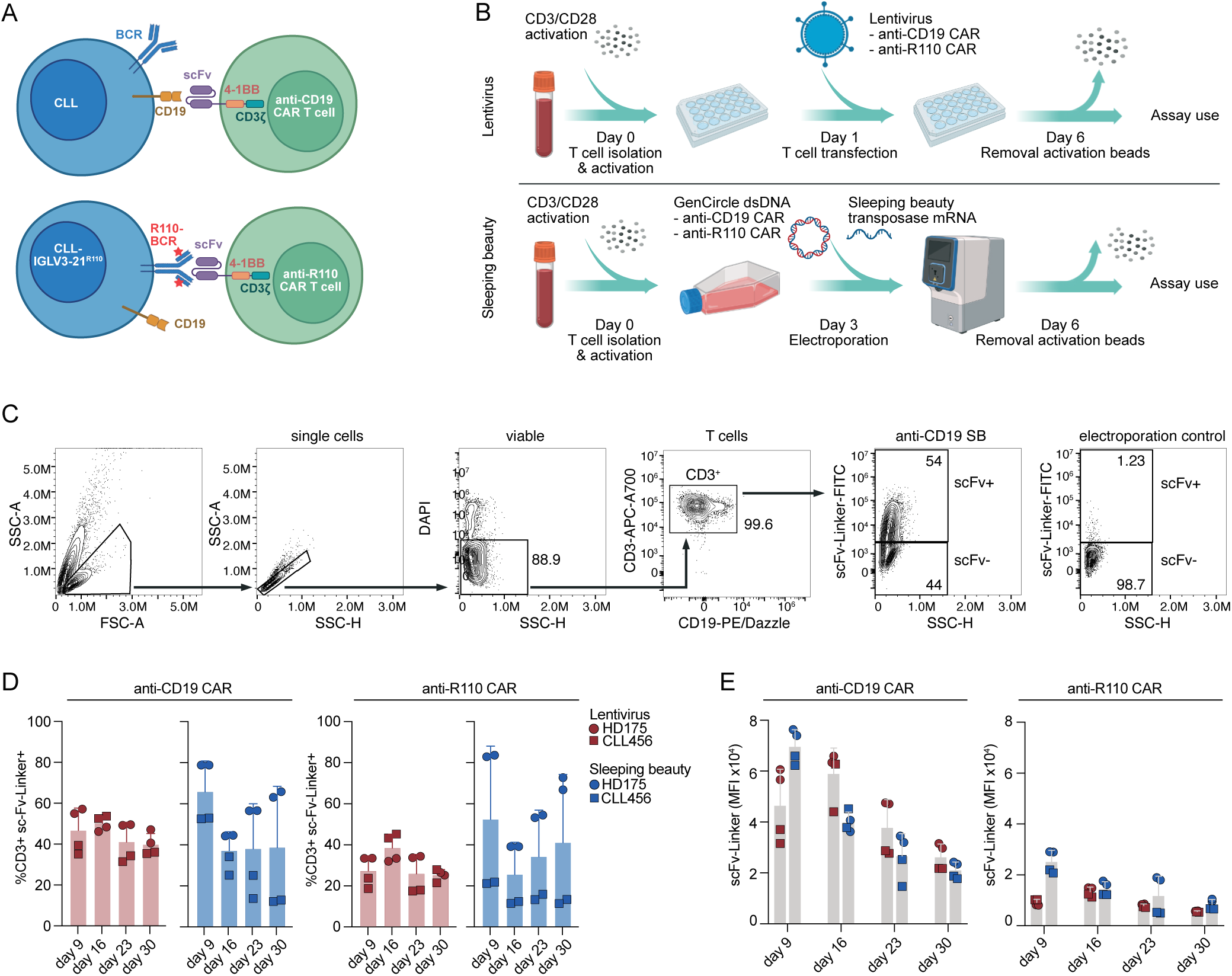
Development of a chimeric antigen receptor (CAR) T cell targeting either CD19 or the IGLV3-21^R110^ neoepitope by LV transfection and SB transposition. **(A)** Schematic representation of the CD19 and IGLV3-21^R110^ (R110) CAR T targeting principle against chronic lymphocytic leukemia (CLL) cells. BCR, B cell receptor. scFv, single-chain variable fragment. **(B)** Schematic representation of the CAR T cell production workflow by LV transfection and SB transposition. **(C)** Gating strategy to determine CAR expression on T cells by flow cytometry. **(D)** Percentage of CAR-positive T cells determined by scFv expression using flow cytometry over the course of four weeks. **(E)** CAR density on T cells determined by the mean fluorescence intensity (MFI) of the scFv using flow cytometry over the course of four weeks. Bars represent the mean ± SD of pooled samples of healthy donor HD175 and CLL patient CLL456 measured in duplicates.

### CD4/CD8 distribution and differentiation stages of anti-CD19 and anti-R110 CAR T cells according to mode of production

Equal production media were used for LV- and SB-manufacturing. These contained IL-15 to maintain less differentiated subpopulations because these are believed to better persist in patients (21, 22). While LV-manufacturing resulted in a CD4^+^ dominant product, SB- manufacturing showed a shift towards CD8^+^ T cells upon expansion consistent with a previous study (Figure 2A and B) (23). The CD8 shift was more pronounced with anti-R110 CAR T cells.

**Figure 2.**
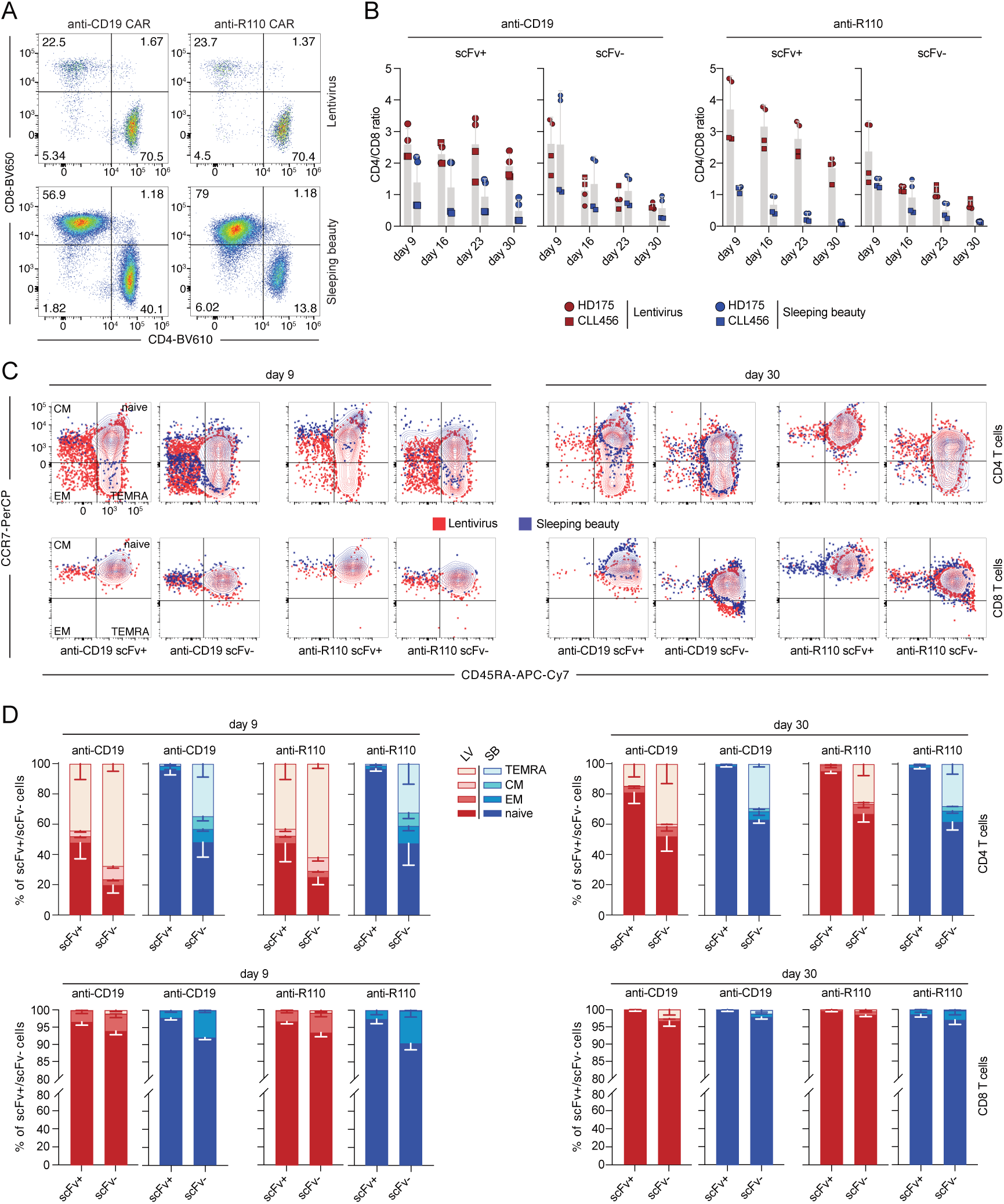
Characterization of the CD4 to CD8 ratio and T cell differentiation of CD19 and R110 targeting CAR T cells. **(A)** Representative image of the distribution of CD4^+^ and CD8^+^ CAR T cells on day 16. **(B)** Calculated CD4^+^ to CD8^+^ ratio of scFv^+/-^ CAR T cells over the course of four weeks. **(C)** Representative T cell differentiation pattern of scFv^+/-^ and CD4^+^/CD8^+^ CAR T cells derived from HD175 measured in technical duplicates by flow cytometry after 9 and 30 days. **(D)** Summarized results of the T cell differentiation pattern of pooled scFv^+/-^ and CD4^+^/CD8^+^ CAR T cell samples of HD175 and CLL456, separated into CD4^+^ and CD8^+^ T cells and according to their scFv expression.

LV- and SB-production modes showed differences in T cell differentiation states of CAR products derived from the same T cell source (Figure 2C). While SB-manufactured products consisted essentially of a naïve-like T cell phenotype (CD45RA^+^, CCR7^+^), LV- manufactured CAR T cells showed a spectrum of CD4^+^ differentiation stages including terminally differentiated cells (CD45RA^+^, CCR7^-^) (Figure 2C). Overall, scFv^-^ T cells better retained the composition of T cell subtypes over time. Figure 2D shows integrated statistics of differentiation stages in T cells derived from a healthy donor and a CLL patient.

### Tonic CAR signaling and co-inhibition in SB- and LV-manufactured products

Interestingly, we observed that in the absence of target cells or antigen, scFv^+^ CD4^+^ T cells manufactured with SB showed an increase of the early activation marker CD69 compared to LV-generated CAR T cells as an indication of tonic CAR signaling (Figure 3A). This was more consistently detected for the R110-CAR T cell product. An increase of CD69 was also observed in scFv^+^ CD8^+^ cells regardless of the manufacturing mode.

**Figure 3.**
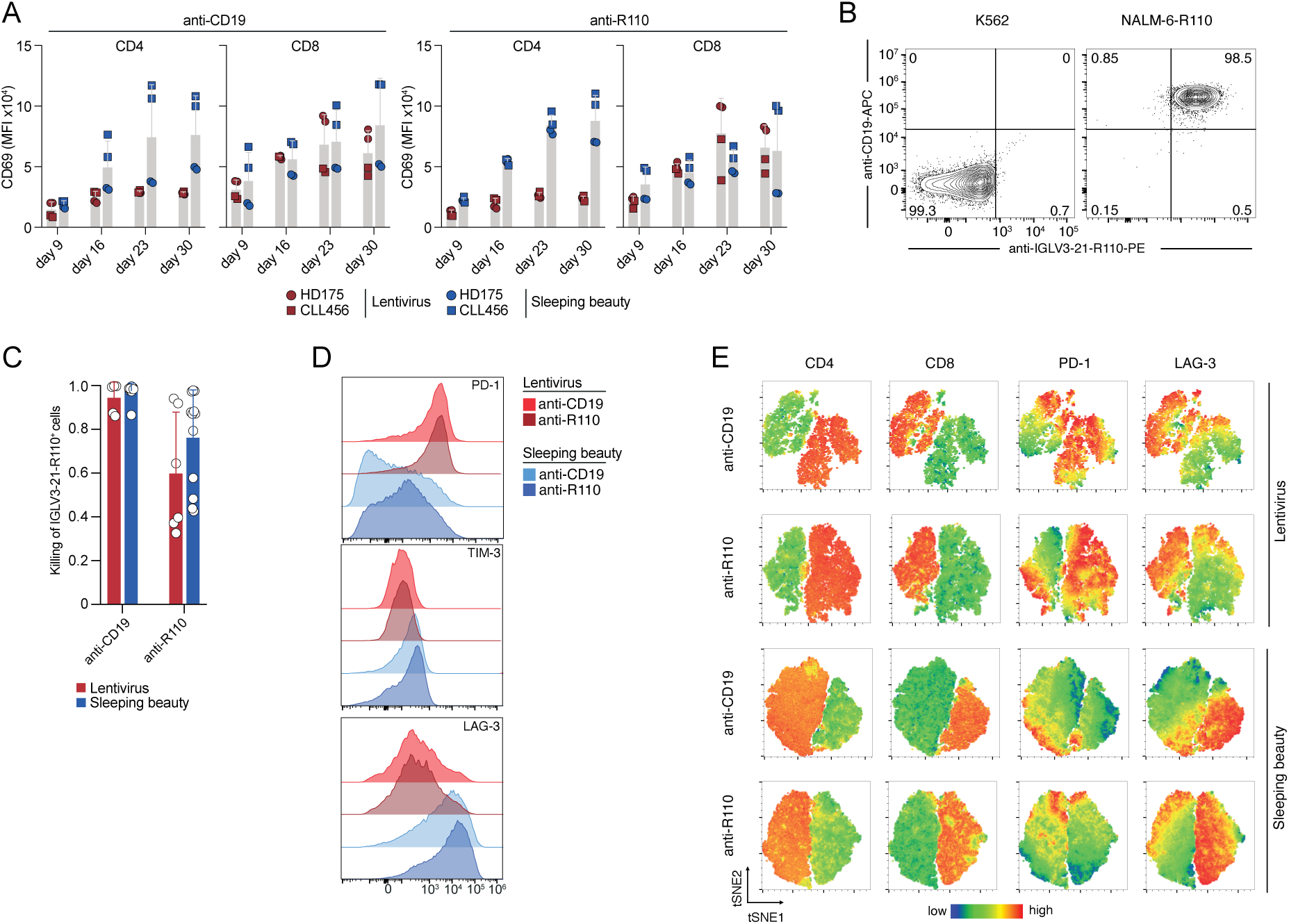
Expression patterns of activation and co-inhibition markers in healthy donor and CLL patient-derived CAR T cells by flow cytometry. **(A)** CD69 surface expression in either scFv^+^,CD4^+^ or scFv^+^,CD8^+^ CAR T cells over the course of four weeks. Bars represent the mean ± SD of pooled samples of HD175 and CLL456 measured in duplicates. **(B)** Representative flow cytometry image of target epitope expression in K562 and NALM-6-R110 cells. **(C)** Pooled results of in vitro coculture assays indicating the killing of CD19^+^, IGLV3-21^R110+^ NALM-6-R110 target cells in relation to CD19^-^,IGLV3-21^R110-^ K562 non-target cells. Bars represent the mean ± SD of pooled experiments. **(D)** Histogram of the PD-1, TIM-3 and LAG-3 expression in CD3^+^ T cells (HD175) on day 9. **(E)** Visualization of the CD4, CD8, PD-1 and LAG-3 distribution pattern of CD3^+^ T cells (HD175) on day 16. Technical duplicate measurements were concatenated and tSNE analysis performed with standard settings.

Next, we asked whether tonic CAR signaling leads to killing of bystander antigen-negative cells. We, therefore, calculated the ratio of specific killing by co-culturing different batches of R110- and CD19-CAR T cells with the antigen-positive NALM-6 cell line transduced with the LeGO-iZeo2 lentiviral vector encoding IGLV3-21^R110^ (NALM-6-R110) and the K562 cell line which is negative for CD19 and the R110 neoepitope (Figure 3B). R110-CAR T cells showed somewhat lower rates of specific killing with more background killing of antigen-negative cells compared to CD19 CAR T cells but this was not associated with production mode (Figure 3C).

Next, we explored whether antigen-independent CD69 upregulation was accompanied by expression of other activation or co-inhibition markers. Therefore, we recorded expression of the classical T cell exhaustion markers PD-1, TIM-3 and LAG-3 in CAR T cells that were cultured without exposure to antigen or target cell line. Measurements were performed between day 9 and day 30 in order to follow dynamics over time. Different marker expression was observed depending on the construct and manufacturing mode as shown in Figure 3D and Supplementary Figure 1. While SB-manufactured products expressed slightly more TIM-3, there were more pronounced differences in PD-1 and LAG-3 expression between production modes. SB-manufactured T cells showed considerably lower PD-1 expression while exhibiting high levels of LAG-3 without antigen encounter. In contrast, LV-manufactured products showed higher PD-1 levels, but instead low LAG-3. When visualizing marker expression by tSNE, LAG-3 expression was virtually restricted to CD8^+^ subsets, while PD-1 was more expressed on CD4^+^ cells and especially in the LV- manufactured products (Figure 3E). Overall, SB-manufactured T cells showed the highest LAG-3 expression in almost all CD8^+^ cells while lacking PD-1 expression.

### Single-cell RNA sequencing of SB- and LV-produced CAR T cells

To investigate the impact of CAR T cell manufacturing methods on the cellular states of engineered T cells, we performed integrated single-cell RNA and single-cell TCR sequencing on CD3-sorted CAR T cell cultures. These cultures included R110- and CD19- CAR T cells produced via SB and LV systems from both a healthy donor and a CLL patient. Untransduced T cells (UTDs) from SB production served as controls. Samples were collected on day 21 of production, yielding a total recovery of 67,799 cells (mean per sample 6,780; range 4,102-8,847) that passed quality thresholds, covering the major T cell subsets (CD4^+^, CD8^+^, and γδ T cells; Figure 4A, B, C). Using standard surface markers and references from published single-cell atlases, we identified a core CD4^+^ cluster with subsets displaying naïve (*CD27*, *CCR7*, *SELL*, *TCF7*), regulatory (*IL2RA*, *FOXP3*, *RTKN2*), and memory-like (*ITGB1*) signatures (Figure 4A, B). A proliferation-associated cluster included both CD4^+^ and naïve-like CD8^+^ T cells (*MKI67*, *SOX4*). CD4^+^ and CD8^+^ T cells were further classified into central memory-like and effector memory-like subsets based on markers such as *CD27*, *GZMK*, *GZMB*, *KLRF1*, *CCR7*, *SELL*, and *LEF1* (Figure 4A, B). In addition, we identified a cluster of memory-like CD4^+^ T cells with T regulatory cell (T_reg_)/effector-like features (*FOXP3*, *KLRB1*, *ITGB1*), including cells expressing the MAIT cell marker gene *SLC4A10*. We also detected γδ T cells with both naïve-like and effector-like characteristics (*TRDC*, *TRGC1*, *GZMB*, *GZMK*; Figure 4A, B). While all clusters were found with both CAR constructs and production modes, SB-manufactured T cells showed a trend towards expanded proliferative CD4^+^/CD8^+^ γδ T cells with high expression of *KLRF1* (proliferation cluster 2, Figure 4A, B). Additionally, γδ T cells and CD8^+^ T cells were more prevalent in the SB product, while the LV product contained a higher proportion of CD4^+^ T cells consistent with flow cytometry (Figure 4C).

**Figure 4.**
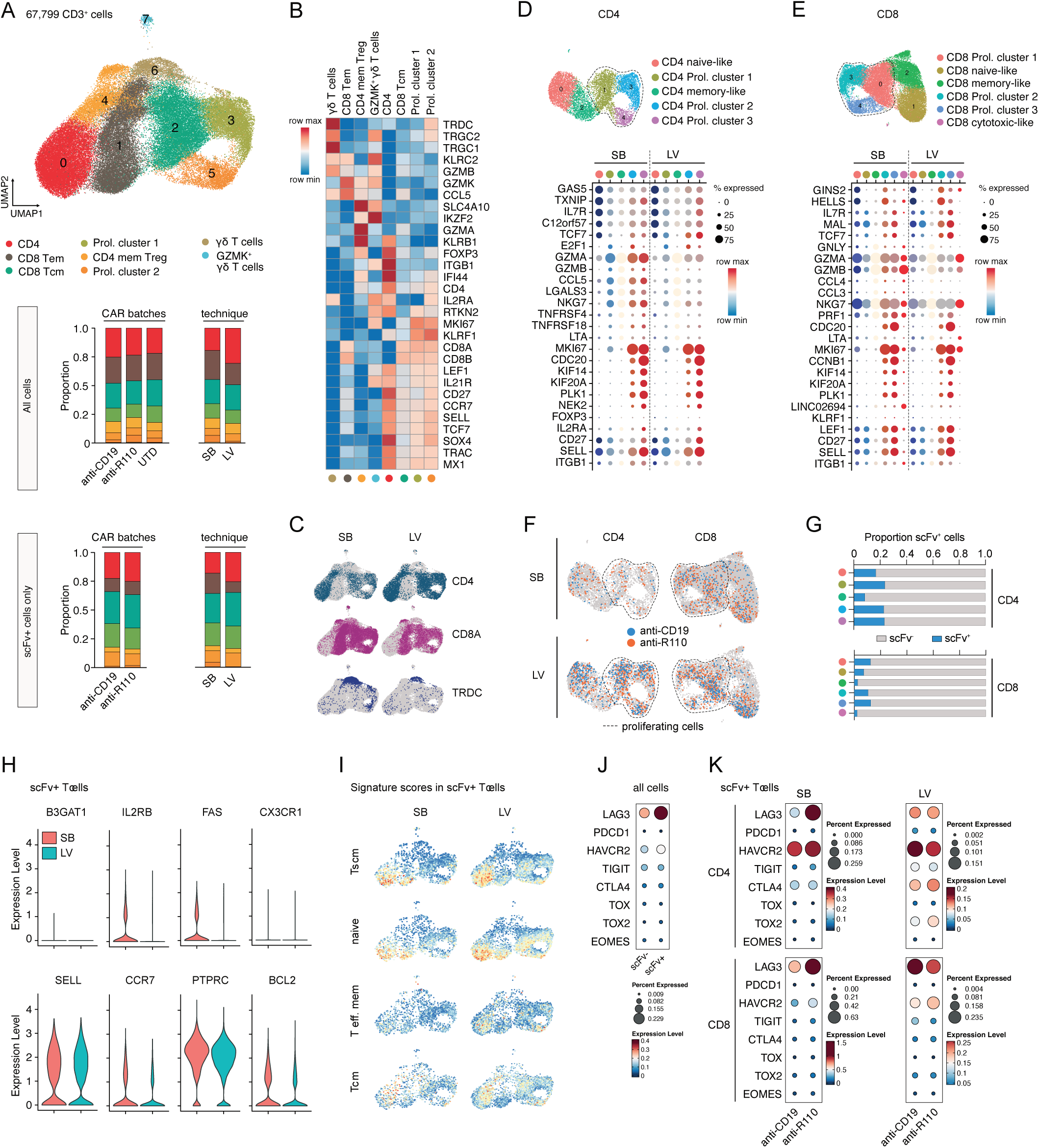
Single-cell RNA sequencing analysis of CD3^+^ T cells from SB and LV production pools. **(A)** Uniform manifold and approximation projection (UMAP) plot of CD3^+^ T cells from the SB and LV production pools. Stacked bar plots display the cluster cell distribution based on CAR T production batches (including UTD control cells) or production mode (SB: sleeping beauty transposition; LV: lentiviral transduction) for all CD3^+^ T cells and CAR T cells only (scFv^+^ cells). **(B)** Heatmap of average normalized expression for conventional T cell marker genes in each cluster. **(C)** UMAP plot showing *CD4*, *CD8A* and *TRDC* expression in CD3^+^ T cells with respect to CAR T production mode. **(D)** UMAP plot of CD4^+^ T cells after re-clustering. Expression of T cell markers and selected cluster-defining genes as dot plot grouped by production mode. Dotted lines indicate proliferating T cell clusters. **(E)** UMAP plot of CD8^+^ T cells after re-clustering. Expression of T cell markers and selected cluster-defining genes as dot plot grouped by production mode. Dotted lines indicate proliferating T cell clusters. **(F)** UMAP plot of CD4^+^ and CD8^+^ T cell subsets with marked scFv^+^ T cells per production mode. **(G)** Proportion of scFv^+^ cells per cluster in (F). **(H)** Violin plots showing expression of marker genes for stem cell-like T memory cells (T_scm_) in scFv^+^ cells from the SB and LV production. **(I)** UMAP plot showing enrichment of gene signature scores for the indicated T cell subpopulations depending on production mode. **(J)** Dot plot showing the expression of selected T cell exhaustion/activation markers in scFv^+/-^ cells. **(K)** Dot plot showing the expression of selected T cell exhaustion/activation markers in scFv^+^ CD4^+^ or CD8^+^ T cells depending on production mode.

Due to overlapping signatures in the overall CD4^+^ and CD8^+^ T cell data, we analyzed these subsets separately after reclustering (Figure 4D, E). Based on conventional T cell markers, cluster-specific differentially expressed genes, and published gene signatures for blood-derived T cells, we identified proliferating cells as well as naïve-like and memory-like subsets within both CD4^+^ and CD8^+^ populations (Figure 4D, E; Supplementary Figure 2A, B). CAR positive T cells expressing scFv were found across CD4^+^ and CD8^+^ clusters, particularly enriched in proliferating and naïve-like subsets, with fewer in memory-like subsets (Figure 4F, G).

In line with the detected CD45RA^+^CCR7^+^ surface patterns (Figure 2C), SB-manufactured CAR T cells exhibited slightly higher expression of markers associated with stem cell-like memory T cells (T_scm_; *FAS* (CD95), *IL2RB*, *CCR7*, *BCL2*), but did not display differences to LV-manufactured cells according to a global established T_scm_ signature (Figure 4H, I). CAR T cells, regardless of production method, showed limited effector memory (T_em_) signatures and some central memory (T_cm_) expression (Figure 4I). In both CD4^+^ and CD8^+^ CAR T cells, co-inhibition markers *LAG3* and *HAVCR2* (TIM-3) were considerably elevated compared to T cells not carrying the CAR (Figure 4J). SB-manufactured R110 CAR T cells showed more *LAG3* and *HAVCR2* transcripts compared to SB-manufactured CD19 CAR T cells (Figure 4K, mind different scales).

### Pathway analysis in SB- and LV-produced CAR T cells

Next, we tested wheter the SB and LV production modes resulted in activation or suppression of specific pathways. For this, we calculated differentially expressed genes (DEGs) in SB- and LV-manufactured products for the CD4^+^ scFv^+^ and CD8^+^ scFv^+^ populations as well as the scFv^-^ bystander cells (Figure 5). Interestingly, across all DEG analyses, scFv^+^ cells consistently yielded higher numbers of DEGs (Figure 5A, D). Independent of scFv expression, CD4^+^ T cells from the SB batch were characterized by increased expression of the chemokine *CCL5* (RANTES), the proliferative transcription factor *HDGLF3* and of cytotoxicity markers (*GZMA* and *GZMB*) including the transcription factor *BHLHE40* that is crucial for regulating T cell effector functions (24) (Figure 5A). Gene set enrichment analysis revealed that T cells from the SB production had higher expression of proinflammatory genes related to TNF, interferon (IFN) and JAK-STAT signaling (Figure 5B). In contrast, SB-manufactured T cells displayed decreased levels of genes for cell cycle control, replication and proliferation (*MYB*, *PCLAF*, *H2AFZ*, *HIST1H1A*) (Figure 5A, B). The scFv^+^ CD4^+^ T cells from the SB production were especially characterized by the enrichment of genes (*DDX60*, *DDX3X*, *TBK1*) that act in the RNA/DNA-sensing RIG-I and TLR signaling pathways including the pivotal mediators and effectors from the NFκB (*IKBKE*, *TRAF2*, *RELA*) and interferon signaling pathways (*IRF3*, *IRF7*, *IFIH1*, *ISG15*) (Figure 5B, C). In addition, CD4^+^ T cells from the SB-batches also had high levels of the E3 ISGylation ligase 5 (*HERC5*) that mediates activating ISGylation of the cytosolic DNA sensor STING (25) (Figure 5A). Notably, we did not observe enriched RIG-I and TLR gene signatures in CD8^+^ CAR T cells from the SB production (Figure 5D, E). Instead, CD8^+^ scFv^+^ T cells were characterized by increased expression of *CCL3*, *CCL4* and the FYN/CSK/Abl interacting transmembrane adatpor protein *PAG1* that negatively regulates T cell activation and is linked to anergy (26, 27) (Figure 5D). Moreover, CD8^+^ scFv^+^ cells displayed enriched p53 pathway signatures, which may be associated with CD8^+^ T cell exhaustion (28), and had increased levels of *LAG3* as compared to cells from the LV production (Figure 5D). CD8^+^ CAR T cells from the SB production shared the general enrichment of pro-inflammatory signatures with their CD4^+^ counterparts (Figure 5D, E). To validate the observed RIG-I/TLR signatures and determine whether they are induced by RNA or DNA in the transfection process, we electroporated CD3^+^ T cells with either the SB mRNA or the CAR DNA constructs alone. Surface marker expression was quantified after 7 days using flow cytometry. The data revealed that, particularly after mRNA delivery, the surface expression of CD69, PD-1, TIM-3, and LAG-3 significantly increased (Figure 5F).

**Figure 5.**
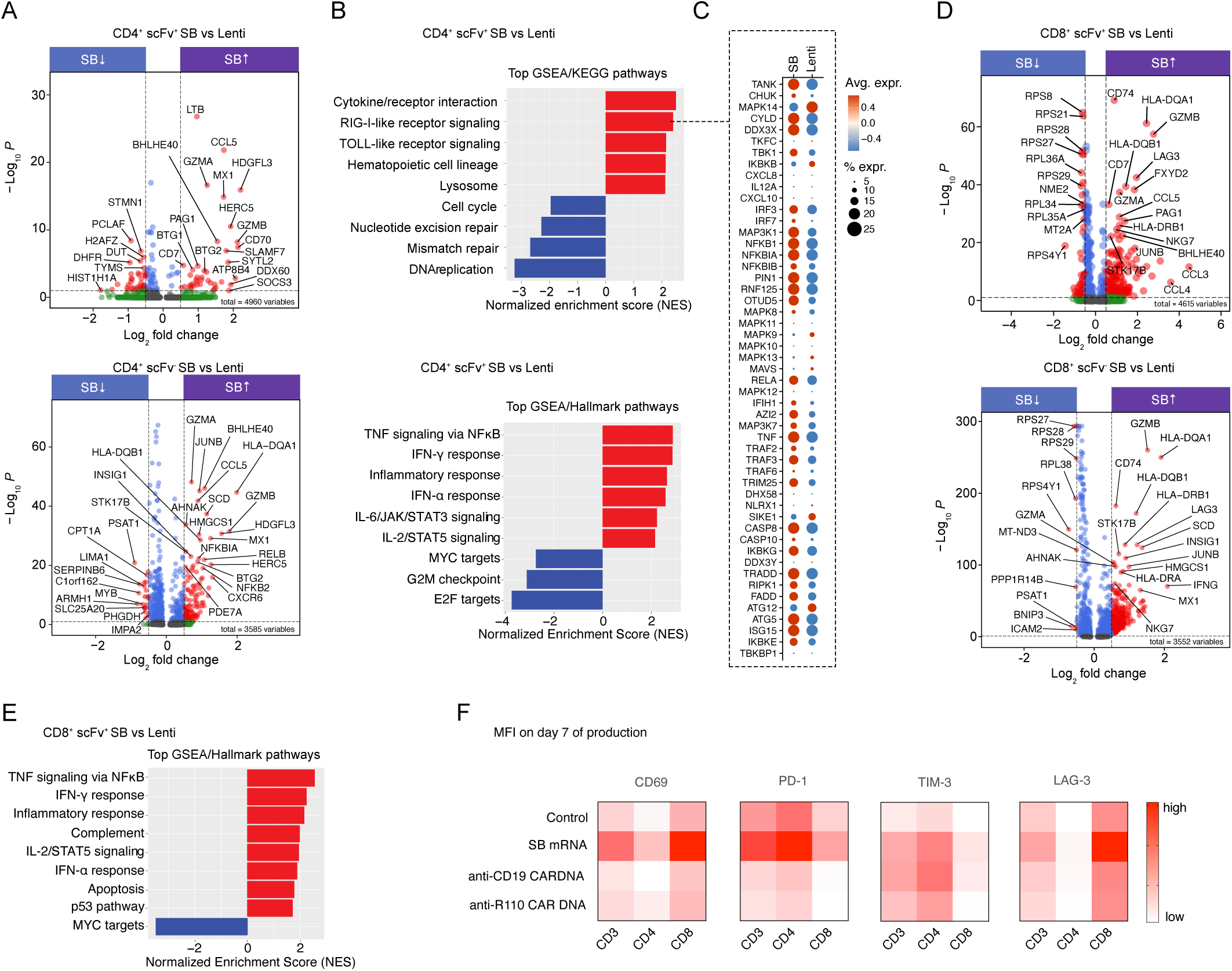
Characterization of production protocol-specific transcriptomic signatures in CD3^+^ T cells from SB and LV production pools. **(A)** Volcano plot showing genes differentially expressed (adjusted P < 0.01) between SB and LV production modes for CD4^+^ scFv^+^ (upper panel) and CD4^+^ scFv^-^ T cells (lower panel). **(B)** Gene set enrichment of GSEA/KEGG (upper panel) and GSEA/Hallmark pathways (lower panel) in SB- manufactured scFv^+^ CD4^+^ T cells as compared to the LV production. **(C)** Dot plot showing the average normalized expression of genes from the RIG-I receptor signaling gene set in SB- and LV-produced scFv^+^ CD4^+^ T cells. **(D)** Volcano plot showing genes differentially expressed (adjusted P < 0.01) between SB and LV production modes for CD8^+^ scFv^+^ (upper panel) and CD8^+^ scFv^-^ T cells (lower panel). **(E)** Enrichment of GSEA/Hallmark pathway genes in SB-manufactured scFv^+^ CD8^+^ T cells as compared to the LV production. **(F)** Heatmap displaying surface expression of indicated markers as MFI on day 7 for mono-electroporation of SB mRNA, anti-CD19 or anti-R110 CAR DNA constructs. Control T cells underwent electroporation only.

### Donor cell constitution impacts transcriptomic signatures of CAR T cells

To test for donor-dependent transcriptome differences, we separated the single-cell data set into HD- and CLL-derived cells (Figure 6A, C). While all defined T cell subpopulations (Figure 4) encompassed cells from both donors (Figure 6A, B), the CLL-derived T cell pools had lower numbers of proliferating cells, but higer numbers of γδ, CD4^+^, CD8^+^ T_em_ and CD4^+^ memory T cells (Figure 6A, B). CLL-derived products had generally increased transcipt levels of *LAG3*, *CD69*, *ID2*, *SLAMF7* several chemokines and lncRNAs as well as HLA class molecules and cytotoxicity markers (Figure 6A, C). In contrast, genes related to cell cycle control and proliferation exhibited pervasively lower transcript levels as compared to HD-derived T cells (Figure 6A, C). Within the reclustered CD4^+^ and CD8^+^ subsets (Figure 4D, E) we detected an enrichment of memory-like cells in CLL-derived T cells independent of production mode, while HD-derived T cells were more enriched for proliferating cells (Figure 6D).

**Figure 6.**
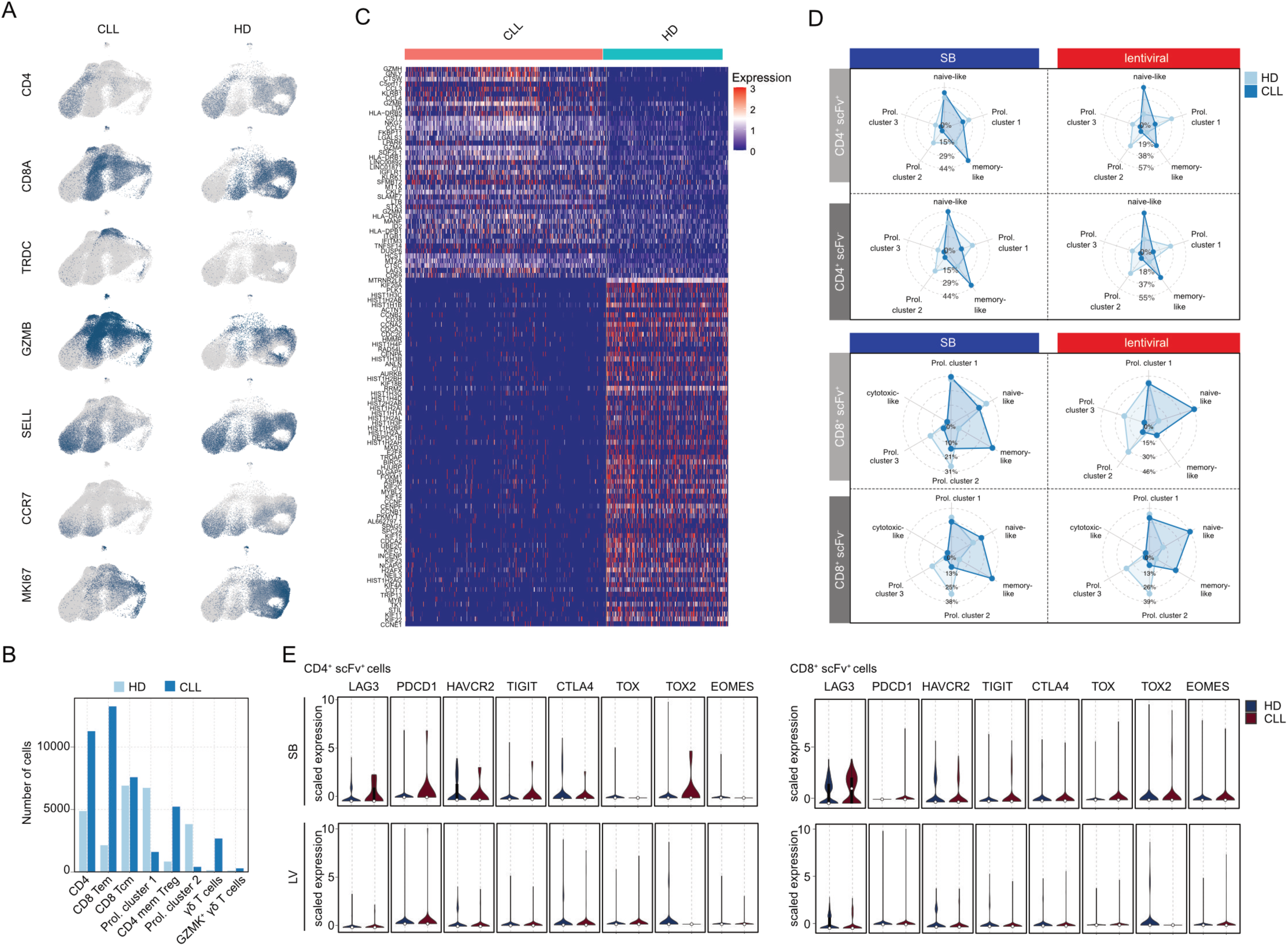
Analysis of donor-dependent impact on SB- and LV-manufactured CAR T cell transcriptomes. **(A)** UMAP plot of healthy donor (HD)- or CLL patient-derived CD3^+^ T cells displaying expression of selected T cell markers. **(B)** Cell numbers in the indicated subpopulations (s. Figure 4) depending on the donor. **(C)** Heatmap showing differential expression of genes between HD-/CLL-derived CD3^+^ T cells. **(D)** Radar plot displaying the contribution of HD-/CLL-derived CD4^+^ scFv^+/-^ and CD8^+^ scFv^+/-^ T cells to the single subsets in the bulk CD4^+^ and CD8^+^ populations from the SB and LV production pools. **(E)** Violin plot showing expression of indicated exhaustion/co-inhibition markers in HD-/CLL- derived CD4^+^ and CD8^+^ scFv^+^ T cells.

Finally, we tested for donor-dependent differences in activation/co-inhibition markers. As compared to HD-derived cells, the SB-associated increase of *LAG3* expression was more pronounced in CD4^+^ than in CD8^+^ scFv^+^ T cells derived from CLL patients (Figure 6E). In addition, we observed higher levels of *PDCD1*, *TIGIT* and *TOX2* in CLL-derived CD4^+^ T cells with scFv expression from the SB production (Figure 6E). CAR T cells from the LV production had lower levels of these markers and showed no clear donor-dependent differences (Figure 6E).

### TCR repertoire analysis of CAR T cells

Single-cell TCR analysis demonstrated polyclonality across all products, with no notable differences between CAR T cells manufactured via SB and LV methods. T cell characteristics, however, varied significantly with the donor source. CAR T products derived from the CLL donor exhibited lower T cell richness and diversity compared to those from the HD, highlighting a restriction in the CLL donor’s T cell repertoire (Figure 7A). This limitation was further confirmed by TCR repertoire overlap analysis, which revealed substantial clonal overlap between the CLL donor’s CAR T products, in contrast to the HD-derived products, which showed minimal clonal overlap (Figure 7B). TRA and TRB gene analyses identified a broad spectrum of gene usage in both donors, with some patterns specific to each donor (Figure 7C). This trend was reflected in a principal component analysis (PCA) of TRBVJ gene usage, where CAR T cell products from each donor clustered together. No distinct differences were observed based on production method or CAR binding moiety (Figure 7D).

**Figure 7.** T cell receptor repertoire analysis of SB- and LV-manufactured CAR T cells. **(A)** Immune metrics. **(B)** Shared CDR3 sequences between donors and production batches. **(C)** Frequencies of TRBV gene usage. **(D)** PCA of TRBVJ usage.

## Discussion

T cell dysfunction is one of the major roadblocks in the development and application of CAR T cell therapies (7). There is only scarce data on how the production mode and the binding moiety may impact CAR T cell phenotypes. Here, we provide a head-to-head comparison of different production modes – lentiviral transfection and sleeping beauty transposition – of R110- and CD19-directed CAR T cells with a focus on T cell activation and co-inhibition marker expression.

In our study, we demonstrate that different CAR constructs exhibit distinct patterns of marker expression for T cell activation and co-inhibition. Notably, we observed substantial differences between SB and LV production methods, even when using identical CAR sequences, T cell donors, and culture conditions – factors known to critically impact the final CAR T cell product (21). Our data suggest that especially the delivery of SB mRNA is a dominant driver of the exhausted-like phenotypes by eliciting intracellular nucleic acid-sensing pathways like RIG-I. Interestingly, upregulation of RIG-I signaling has been reported in terminally exhausted CD8^+^ T cells and was linked to reduced CD8^+^ T cell effector functions in tumor-infiltrating lymphocytes (29). Also, other product-defining features were significantly affected by the production mode, such as CD4/8 ratio, differentiation stage or CD69 expression as a marker for antigen-independent tonic activation. These findings are crucial as they suggest that the optimization of CAR constructs extends beyond the design of co-stimulatory domains or the configuration of the binding moiety supporting that each construct must also be rigorously tested under various manufacturing conditions.

In a related approach, a previous study compared the phenotypes of CD19-targeting CAR T cells either manufactured using electroporation and the PiggyBac transposase system or produced by lentiviral transduction (30). The authors noted higher CAR-density and cytokine release in CAR T cells generated with the transposase system, but no significant differences regarding CD69 or PD-1 surface expression or anti-tumor efficacy in a murine xenograft model (30). CAR T cells produced with the transposase system displayed lower frequencies of T_cm_ and T_em_ phenotypes as their lentiviral counterparts and had higher expression of genes related to immunological and inflammatory pathways as shown in our study (30). However, their analyses were restricted to the first week after CAR T production (30).

There is some data linking CAR T cell phenotypes to efficacy outcomes of CAR T cells in patients. While some studies suggested that a balanced ratio of CD4 and CD8 cells in the product would support persistence and eradication of the malignant clone in patients (31, 32), others did not support this conclusion (8). Moreover, in one study early memory T cell differentiation in the CAR T cell product was found to correlate with better responses while an effector or exhausted T cell differentiation was not (8). In another study, peak CAR T cell detection in patients correlated with the percentage of a naïve CAR^+^ CD8^+^CD45RA^+^CCR7^+^ T cell subset in the CAR T cell product (33). Furthermore, the percentage of PD-1^+^CD8^+^ CAR T cells coexpressing either TIM-3 or LAG-3 correlated with a lower response to therapy (8). CD62L (*SELL*), also known as L-selectin, is another marker of naïve T cells, including stem cell memory T cells, but also central memory T cells. Its enhanced expression was shown to lead to better control of tumor growth in an in vivo model and its expression level was associated with a better ex vivo expansion (34–36). Overall, T cell subset distribution and co-inhibitory receptor expression as well as prior antigen encounter influences outcomes with CAR T cells (21, 36, 37), suggesting that the differences associated with production mode and CAR specificity seen in our study may be of clinical relevance.

The determinants of T cell dysfunction in the setting of CAR T cells are still rather ill defined. T cell exhaustion is considered the result of continuous exposure to antigen (7, 38) but may also occur without antigen exposure in the case of CAR T cells, e.g. by self-aggregation of the CAR resulting in tonic signaling (7). It has been assumed that tonic signaling could not only be a feature of the CAR sequence itself, but also a function of CAR membrane density. To this end, “self-driving” armored CAR T cells that upregulate CAR expression only in case of antigen encounter were developed and tested preclinically (39). As opposed to this assumption, we found that the pattern of high LAG-3 expression was not associated with higher expression of CAR molecules. In contrast, we observed higher basal LAG-3 expression in SB-manufactured R110-CAR T cells that showed much lower CAR density than CD19-directed CAR T cells. This suggests that – at least in our model – high CAR expression is not a critical inducer of tonic CAR signaling.

We observed a low percentage of scFv^+^ T cells and a depletion of proliferative T cell clusters in CAR T cells derived from the CLL donor. These effects were more pronounced in SB-manufactured products, suggesting that the stress associated with SB transposition may further impact the already fragile T cells from patients, thereby impairing their proliferative capacity. In conditions like CLL, where T cell defects are well-documented, it will be essential to investigate this further to optimize transgene delivery and achieve a cell product that can proliferate and persist effectively in patients.

In summary, while CAR T cells hold great promise in oncology, T cell functionality remains a critical challenge, particularly in the autologous setting. Addressing this issue is essential for enhancing the efficacy and expanding the applicability of CAR T cell therapies across a broader range of cancers.

## Acknowledgements

This study was partially funded by SinABiomedics as well as the Mertelsmann Foundation, the innovation focus cell therapies at the University Hospital of Base and the Swiss National Science Foundation (SNSF) [10.001.762] (all to M.B.).

## Author contributions

Idea & design of research project: MB; Supply of critical material (e.g. patient material, cohorts): MB; Establishment of Methods: SS, CS, PBS, AZ, CF, HL; Experimental work: SS, CS, PSB, CF, AZ, NF; Analysis and interpretation of primary data: MB, CS, SS, SA, PBS; Drafting of manuscript: MB, CS, SS. All authors reviewed and revised the manuscript.

## Code availability

No code has been developed for this study.

---

**Supplementary Table 1.**
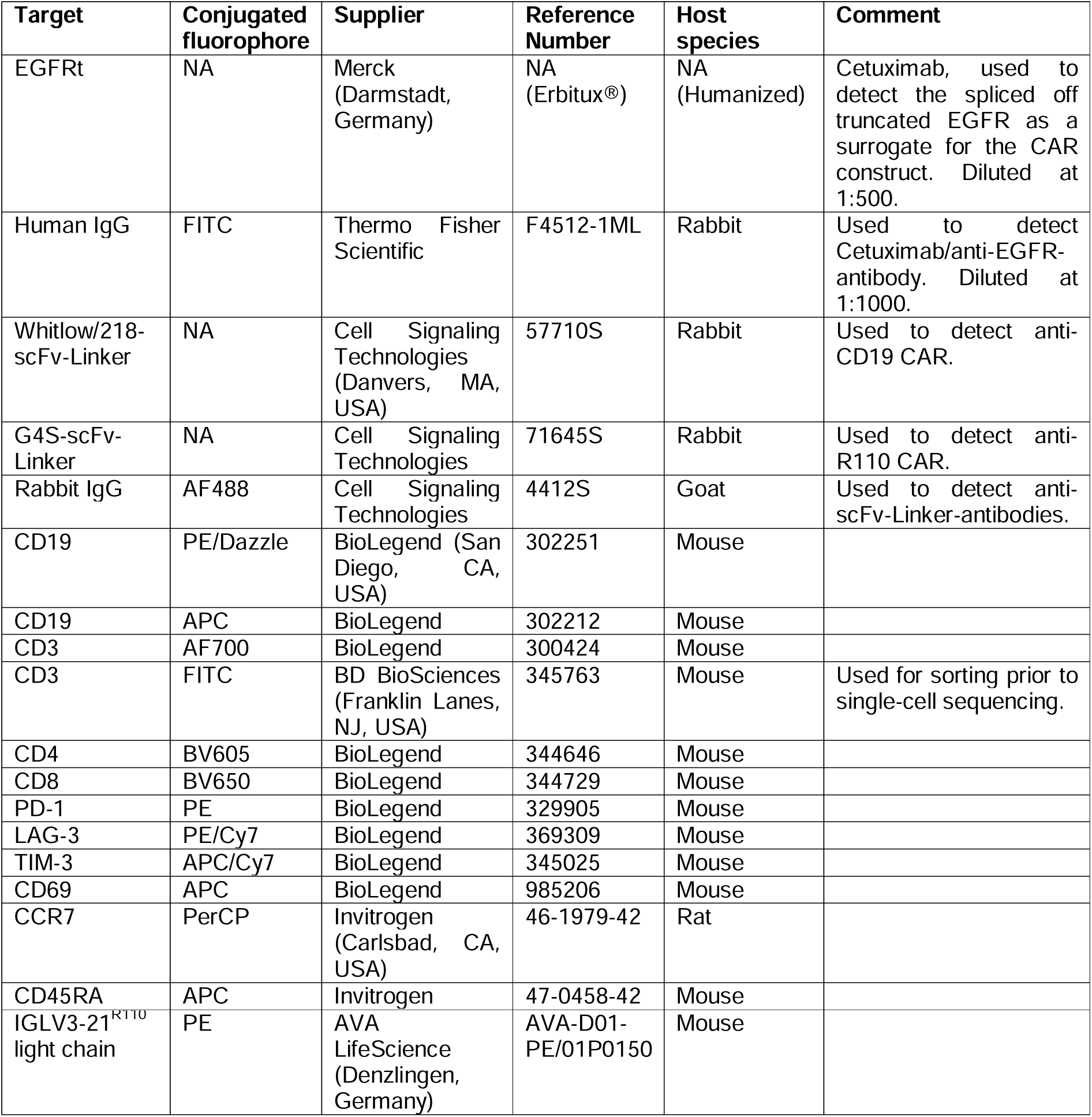
Antibodies used for flow cytometry.

**Supplementary Figure 1. Expression patterns of activation and co-inhibition markers in healthy donor and CLL patient-derived CAR T cells by multicolor flow cytometric analysis.(A)** CD69, **(B)** TIM-3, **(C)** LAG-3 and **(D)** PD-1 mean fluorescence intensities (MFI) in CAR T cells separated by CD4, CD8, scFv^+^ and scFv^-^ expression. Bars represent the mean ± SD of pooled samples of HD175 and CLL456 measured in duplicates.

**Supplementary Figure 2. Top 10 differentially expressed genes per cluster in the CD4^+^ and CD8^+^ subsets.**

