## Supplementary figures and images for "CAR T cell engineering impacts antigen-independent activation and co-inhibition"

### Supplemental Figure 1

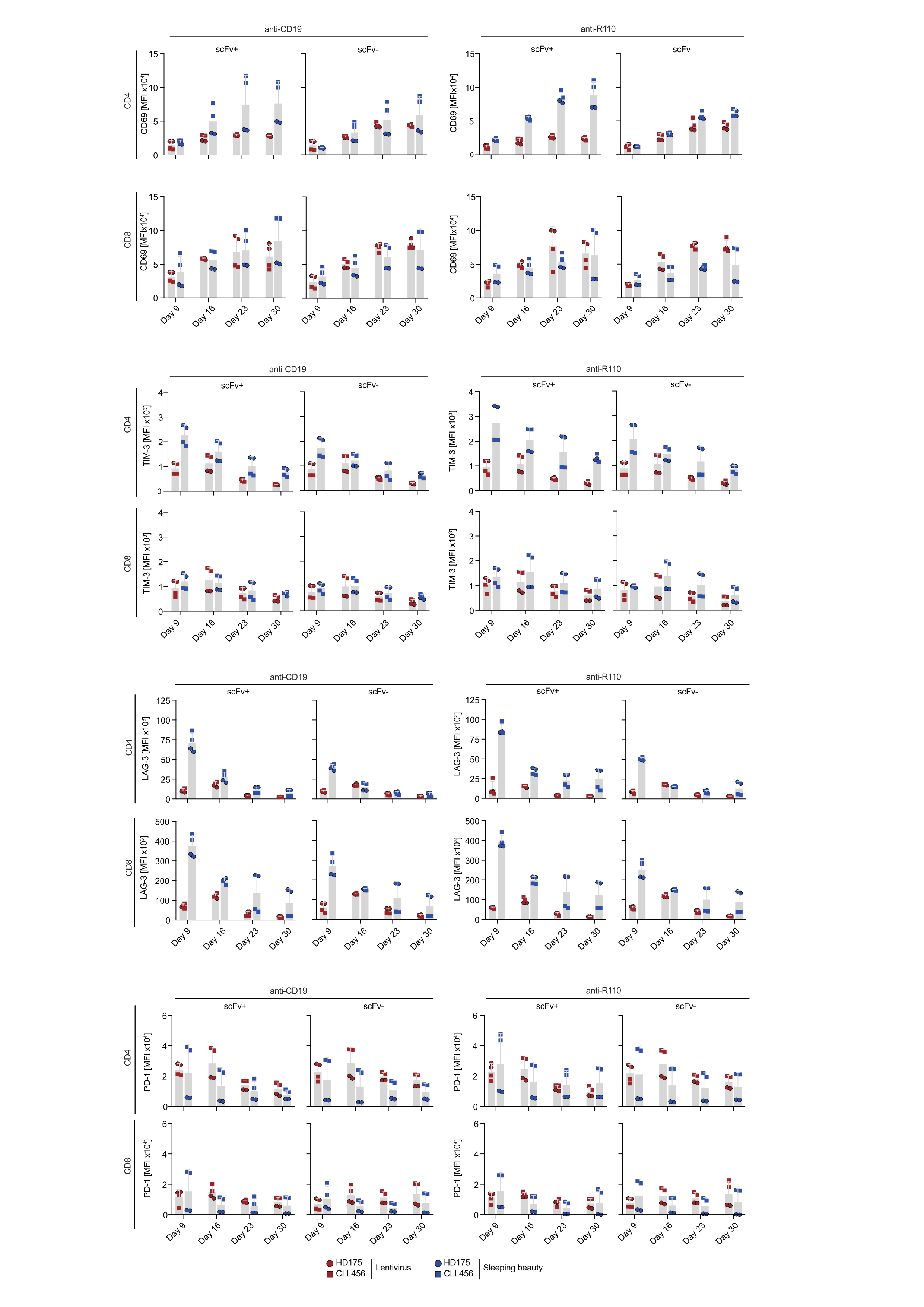

### Supplemental Figure 2

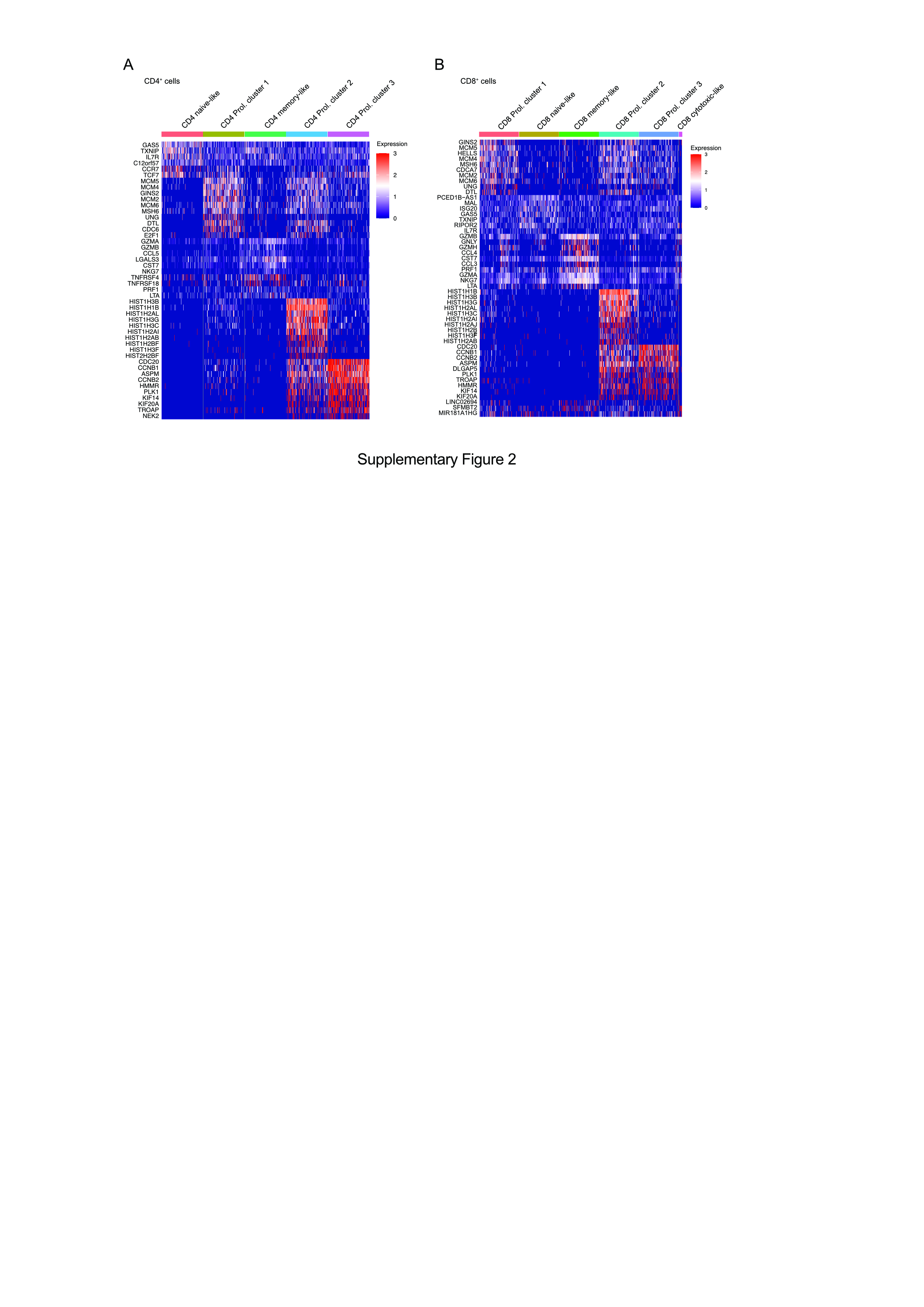
